# Factors influencing *Agrobacterium*-mediated genetic transformation efficiency in wheat

**DOI:** 10.1101/2023.12.01.569549

**Authors:** Thiyagarajan Karthikeyan, Noguera Luis Miguel, Mario Pacheco, Velu Govindan, Prashant Vikram

## Abstract

Assessment of the efficiency of *Agrobacterium tumefaciens* mediated transformation in four bread wheat varieties was focused and upon optimization of experiments done to obtain tangible results. The four varieties namely Fielder, Navojoa, Baj and Kachu/saul were transformed with optimized protocols, subsequently randomly selected from selection medium amalgamated with phosphinothricin (Glufosinate). Initially the variety Fielder was proceeded with 36 different concentrations of synthetic auxins 2,4-D and Picloram as 36 treatments. The treatment 30 (48 μg of 2,4-D and 64 μg of Picloram) has shown higher efficiency and is comparable in terms of the three traits studied. Additionally, antioxidants and growth regulators were adjusted to obtain better result towards reasonable transformation frequency. Based on the optimization for the variety Fielder, additional varieties were also tested and selected against the selectable marker PPT. Selected genotypes were subjected for *gusA* or *Bar* gene PCR amplification. The result revealed the frequency of transgenesis with the range of 60-70% with the varieties studied, however the whole experiment needs further repetitions for confirmatory result. Transgenic genotypes of the Fielder variety showed moderate to strong *GUS* expression in leaves, anthers, seeds, seed coats, roots compare to non-transgenic control plant tissues. PPT leaf painting assay showed the lack of necrosis on painted area of leaves in transgenic genotypes compared to the non-transgenic ones.

## Introduction

Successful genetic transformation through *Agrobacterium* tumefaciens relies on several factors, including host specificity. Earlier, it was believed that *Agrobacterium* transformation was specific to dicots (Songstad et al., 1995) rather than monocots, but several improved protocols in monocots led to the feasibility of *Agrobacterium*-mediated transformation in Gramineae, including major cereal species such as wheat (Cheng et al., 1997; He et al., 2010), rice (Ozawa et al., 2012), and maize (Ishida et al., 2007). Although *Agrobacterium*-mediated transformation was successful in a few varieties of wheat, such as Bobwhite, other varieties required extra virulent genes in the helper plasmid for successful transformation of the T-DNA-integrating plasmid; hence, the plasmid pSoup was manipulated and constructed with an additional extra virulence gene called *vir G* to enhance the transformation in extended wheat varieties (Amoah et al., 2001; Ke et al., 2002). Otherwise, plasmids such as pAl154, pAL186, or pTOK233 containing a 15-kb Komari fragment carrying a fixed sequence of virulent genes such as *vir B, C*, and *G* were used to enhance the efficiency of transformation in wheat (Amoah et al., 2001; Wu et al., 2008; He et al., 2010). The biolistic method is an alternative approach to *Agrobacterium*-mediated genetic transformation in plants; the majority of the wheat transformation in wheat was done using this alternative approach. The advantages of *Agrobacterium*-mediated transformation over other alternative mechanized methods include the delivery and integration of low copies of the gene of interest (quite often one or two), integration of large segments (even up to 10 kb), minimal rearrangements due to defined ends of T-DNA (Ishida et al., 2007), and relatively higher co-expression of the introduced gene in the host with facile in vitro manipulation (Cheng et al., 1997).

Higher transformation efficiency was recently reported in wheat using *Agrobacterium*-mediated transformation, from 51% up to 90% (Richardson et al., 2014; Ishida et al., 2015). Elite cultivars have reportedly been shown to be recalcitrant to genetic transformation, which results in lower transformation efficiency (Luo et al., 2021). However, here we are reporting the efficiency of *Agrobacterium*-mediated transformation from 60% to 70% upon appropriate modifications of protocols. Numerous factors play a vital role in the efficient *Agrobacterium*-mediated transformation, including the size and stage of the undifferentiated tissue explant, efficient strain selection, media compositions, physico-chemical conditions of the media such as pH and hormonal adjustments, and efficient usage of signal transducer acetosyringone, growth promoter phloroglucinol, etc. Undifferentiated tissues are the pivotal source of the explant; therefore, the type (immature embryo) and size (Bartlett et al., 2008) are the key factors that promote the efficiency of *Agrobacterium*-mediated transformation. If the size is less than 1.5 mm, it leads to the lysis of the tissue without further developmental stage progression. Although the timely collection of immature embryos after the 11–16 post-anthesis period (Jones, 2015) is not always expected to provide the correct size, in this case, the vernalization of embryos at 4°C for 3–4 days would substantially increase the desired size of an embryo (up to 2 mm in diameter) without allowing over-maturity.

The transformation of T-DNA from Ti Plasmid to plant cell chromosomes is actively induced by the expression of a native wound-related phenolic compound called acetosyringone (Stachel et al., 1985), which persuades the expression of the *Agrobacterial vir* gene under specific ambient temperatures. The external supply of 200μM acetosyringone in tissue culture media during transformation remarkably increases the T-DNA integration in the host chromosomes, and the successful examination of the silicone-based surfactant Silwet-77 during transformation was also positively evaluated (Wu et al., 2003). An additional compound known as phloroglucinol synergistically acts with auxin by partially inhibiting the effect of cytokinin. Phloroglucinol increases transformation efficiency up to 75% upon combining 25 mg/L of this compound with an appropriate amount of acetosyringone (Blanco et al., 2012). The phytohormone auxin is a vital factor that induces embryogenesis and differentiation (Möller and Weijers, 2009). Two synthetic auxins, namely 2,4-dichlorophenoxyacetic acid (2,4-D) and Picloram, stimulate embryogenesis and subsequent callus regeneration in cereal tissues (Wernicke W., Milkovitz et al., 1987). Adjustment of these compounds in *Agrobacterium*-mediated transformation from inoculation towards co-culture, callus induction, and regeneration plays an essential role in the successful development of transgenic genotypes with higher transformation efficiency. *Triticum aestivum* (Bread wheat or common wheat) is an allohexaploid species, contributing a major staple food supply to the global population along with succeeding crop species rice and maize. Estimated annual wheat production spanning the year 2016-2023 was 732-783 million tons (FAO, 2016-2023); however, global wheat demand would be expected to increase by 60%–110% by 2050 (Tilman et al., 2011). The genetic transformation through *Agrobacterium* in wheat is limited, rather than the amount of transformation conducted through biolistic approaches. During the past decade, specific attention has been given to *Agrobacterium*-mediated transformation with successful expression of the introduced gene. However, the efficiency of the transformation was limited and varied from one genotype or variety to another; thus, the current experiment was conducted to unravel the efficiency of wheat transformation in four wheat cultivars upon modifying or optimizing the protocols.

## Materials and Method

### Transformation procedure

The transformation technique was adapted from Ishida et al. (2015) and subjected to necessary alterations with the ongoing optimization of the CIMMYT protocol based on the necessity for specific genotypes or varieties. About 430 μl of *A. tumefaciens* stock culture was inoculated into 10 mL of MG/L medium containing 20 μl of 100 mg/ mL carbenicillin and 20 μl of 50 mg/ mL Kanamycin and incubated overnight at 37 °C in a shaker at 250 rpm. Bacterial growth was observed with turbidity (OD at 0.4 at A600) the next day, and the culture tubes were centrifuged at 3500 rpm (2000 g) in a Thermo Scientific centrifuge for 10 min. The embryo was extracted from each seed under a light microscope after excising the radicle with the help of a scalpel and forceps. *A. tumefaciens* pellet obtained from centrifugation was briefly mixed with 4800 μl of 1X liquid media containing an appropriate amount of silwet, acetosyringone, and phloroglucinol. Isolated embryos (about 30 embryos for each microfuge tube) were immersed in 600 μl of the 1X inoculation medium with *A. tumefaciens*.

The tubes were mildly vortexed three times for 15-20 seconds at 10-minute intervals and incubated in a water bath for 3 minutes at 46 °C. The infected embryos were transferred to the inoculation/co-cultivation solid medium, which allows the bacterial growth as well as the induction of embryos towards the formation of calluses due to the infection of *A. tumefaciens*. Because of the absence of plant hormones, the induced embryos were not subjected to callus formation or regeneration during the period of co-cultivation and induction. Subsequently, the induced embryos were transferred to the callus regeneration medium, which selectively allows the callus but not *A. tumefaciens* to grow because of the presence of timentin, an antibiotic that selectively acts against the bacterium. The presence of an appropriate amount of synthetic auxin hormones, such as 2,4-D (50 μg = 500 μl of 100 mg/l) and Picloram (50 μg = 500 μl of 100 mg/l) for each petriplate containing 100 mL volume of the medium, allows the formation of callus from each embryo upon incubation for 3–4 weeks at 26 °C in a growth chamber without light.

A further hormone adjustment was made, and 36 different combinations were tested as 36 different treatments. The callus was then transferred to the shoot and root induction medium with appropriate amounts of plant hormones and placed in a light chamber at 26 °C for 4 to 5 weeks. Subsequent selection with 1X WLS medium (for 200 mL) containing phosphinothricin (100 μl of PPT 5 mg/ mL) and 140 μl of 300 mg/ mL Timetin for 4 to 5 weeks (two times) and hardening of the plants was done in soil-filled mini pots (volume: 150 cm^2^) for 4 to 5 weeks.

### *GUS* assay for various tissues

Ingredients: I. 1M Phosphate buffer: 43.55 g of K_2_HPO_4_ and 34.02 g of KH_2_PO_4_ dissolved in 250 mL of distilled water, autoclaved, and stored at 4°C. II. 100 μl Dimethyl Sulfoxide. III. 5 mg X-Gluc. IV. 100 mM Phosphate buffer was prepared from 1M stock buffer using sterile distilled water, and pH was adjusted to 7. Approximately 5 mg of fresh X-Gluc was dissolved in 100 μl of DMSO in a 25 mL conical tube, and 9 mL of 100 mM Phosphate buffer (pH 7) was added in the same tube, mixed thoroughly, and wrapped with aluminum foil to avoid light penetration. It was stored at 4°C for immediate use. This solution must be used within one month from the date of storage to avoid the loss of sensitivity of the X-Gluc. Various tissues, such as leaves, roots, stems, young seeds, the tips of the young spikes, and anthers from transgenic genotypes of varieties Fielder or Navojoa, were collected using multiple 2 mL tubes and kept in containers to avoid the denature of the β-glucouronidase from the expressed tissues. Approximately 300 to 500 μl of X-Gluc buffer (pH 7) was added to each tube containing the tissues and incubated at 37°C for overnight. PPT (Phosphinothricin) leaf painting assay and PCR were carried out as reported with our research communications (Thiyagarajan et al., 2022).

### Statistical analysis

Callus size, callus frequency, and shoot frequency per callus were measured from all 36 treatments with duplication. An analysis of variance (ANOVA) was carried out, and Least Significant Difference (LSD) with Bonferroni correction was estimated using R (R Core Team, 2014).

## Results

### Effects of vernalization and protocol optimization

The number of genotypes examined in these four varieties of bread wheat revealed the presence of an optimal transgenic frequency. The result obtained upon optimizing the procedures of *Agrobacterium*-mediated transformation is as follows: Seeds of the wheat variety Fielder were collected after 14 days (Days After Flowering) and kept at 4 °C for two to three days to achieve optimum growth of embryos without breaking the dormancy of seeds. This optimization resulted in embryos with a size range of 1.5 to 2 mm at the time of excision for infection and transformation. The surface-sterilized embryo was extracted from each seed under a light microscope, and the radicle was excised with the help of a scalpel and forceps.

Isolated embryos (about 30 embryos for each microfuge tube) were transferred to 600 μl of the 1X inoculation medium containing 1% glycerol (as a surface protectant), 9 μl (0.015 μM/μl) of Silwet solution (organosilicone surfactant), and 5 μl of acetosyringone (250 μM) (signal transducer) and phloroglucinol (410 μM) (growth promoter) solution. Further, oxidants (1 mM L-cysteine), adsorbents (0.5 mM PVP), and the control (only inoculation solution) were also tested in separate tubes containing 1X inoculation solution prior to transformation. All these compounds enhanced the efficiency of transformation, while PVP 0.5 mM was not as effective as other compounds.

### Physical stress-based treatments to enhance the *Agrobacterium*-mediated transformation

The *A. tumefaciens* pellet obtained from centrifugation was briefly mixed with 4.8 mL of 1X inoculation solution containing 72 μl of Silwet solution and 40 μl of acetosyringone (250 μM) and phloroglucinol (410 μM) solution. About 600 μl of the solution, along with 430 μl of *A. tumefaciens*, was transferred to a microfuge tube containing approximately 30 embryos. The tubes were mildly vortexed three times for 15-20 seconds at 10-minute intervals and incubated in a water bath for 3 minutes at 46°C (Treatment 1). They were briefly vortexed in order to achieve the bacterial infection on embryos. The infected embryos were transferred to the induction medium, which allowed for bacterial growth as well as the induction of the embryo towards the formation of a callus. This medium doesn’t possess any plant hormones; hence, it allows the induction of embryos towards callus formation for three days. Apart from the first treatment, additional treatments were also given during transformation for different sets of embryos of the same variety. Treatment 2: 30 embryos were centrifuged at 14,000 rpm for 1 minute with *A. tumefaciens*. Treatment 3: 30 embryos were centrifuged at 14,000 rpm for 1 minute without *A. tumefaciens* and further incubated with *A. tumefaciens* at 46°C for 3 minutes. *GUS* assays and callus regeneration efficiency concluded that treatment 1 was more efficient than other treatments during transformation.

### Assessment of transformation efficiency with the *GUS* assay

About 15% of the transformed embryos were subjected to a *GUS* assay with dark chamber incubation at 37 °C for 12 to 24 hours. The observation revealed the efficiency of transformation due to the formation of blue-colored spots along embryos after 12 hours, while further incubation up to 24 hours intensified the blue-colored spots. However, prolonged incubation caused endogenous *GUS* activity; thus, the observation of 12-hour incubated embryos showed optimum transgenic *GUS* expression due to the observation of systematic blue spots, unlike endogenous expression (Fig. 1).

**Figure 1.**
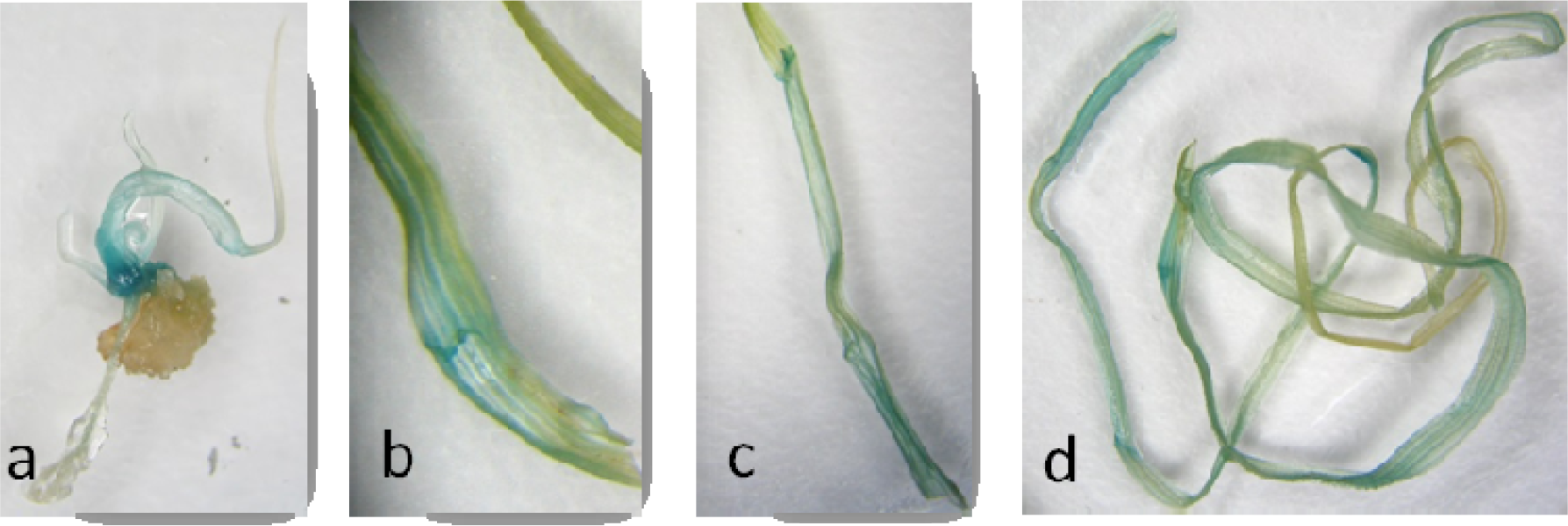
*GUS* expression from the juvenile shoot/leaf tissues formed from the callus for the variety Navojoa.

### Effects of the addition of antioxidants and growth promoters

The infected embryos were transferred to the induction medium, which allowed the bacterial growth as well as the induction of the embryo towards the formation of a callus because of the *A. tumefaciens* infection. Because of the absence of plant hormones, the induced embryos did not undergo callus formation or regeneration during the period of co-cultivation and induction. In this phase, the embryos were allowed to grow for three days in a growth chamber at 26°C without light in order to achieve callus induction. The induced embryos along with adhered *A. tumefaciens* were transferred to callus regeneration medium, which selectively allows the growth of callus alone due to the presence of an antibiotic timentin (Ticarcillin and Clavulanate), which inhibits the growth of gram-negative *A. tumefaciens* by arresting its cell wall synthesis. The presence of an appropriate amount of synthetic auxin hormones, such as 2,4-D and Picloram, allows the formation of a callus in each embryo. Callus regeneration was enriched with 0.5 mM PVP or 1 mM cysteine. For control, the medium was inoculated without the addition of PVP or cysteine. These treatments were replicated three times each. Just prior to the preparation of the medium, 100 mg/l of ascorbic acid, 0.85 mg/l of AgNO_3_, 250 mg/l of Carbenicillin, 100 mg/l of Cefatoxime and 0.4 mg/l of 2,4-D, 0.3 mg/l of Picloram, and 1mM Cysteine were added. In this phase, embryos were allowed for three weeks in order to achieve callus generation without bacterial growth at 26°C in a dark chamber for regeneration of the callus. At the end of the 3rd week, the size of the callus and frequency of callus formation in the regeneration medium were recorded for statistical analysis. During callus regeneration, different concentrations of 2,4-D and Picloram (16 μg/μl, 24 μg/μl, 32 μg/μl, 40 μg/μl, 48 μg/μl, 64 μg/μl) resulted in the combinations of 36 treatments in total. Among which, the combination of 16 μg/μl 2, 4-D with all 6 concentrations of Picloram revealed better callus regeneration (Treatment 1 to 6) and subsequent multiple shoot induction after 3 weeks in a light chamber at 26°C. The size of all calluses from each treatment with a duplicate was measured to calculate the average size of the callus from each treatment (Fig. 2).

**Figure 2.**
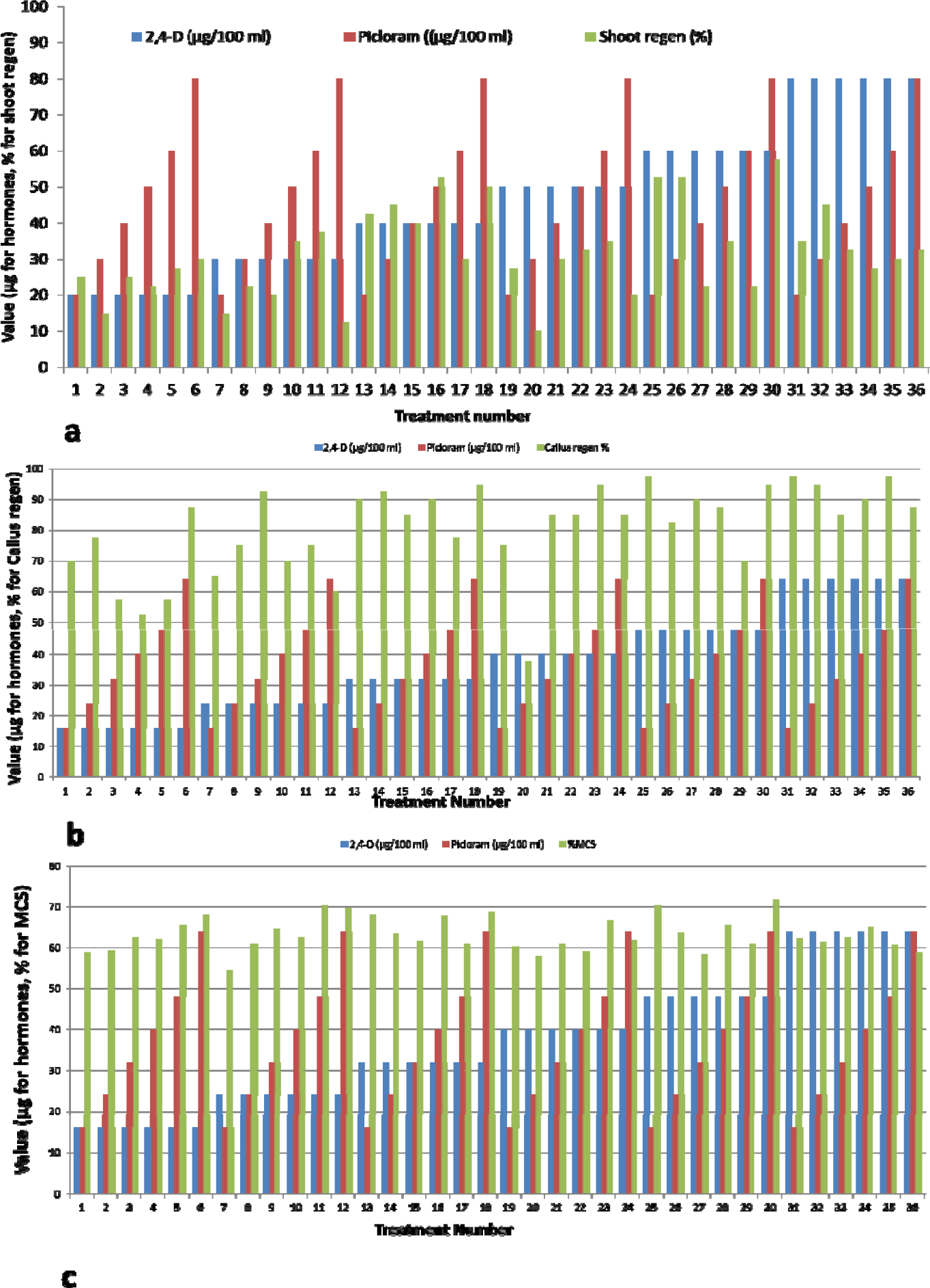
Bar diagram showing 36 treatments combinations of 2,4-D and Picloram and their effect on percentage of shoot regeneration, percentage of callus regeneration and mean callus size.

### Regeneration of shoots and roots

Embryos from all the treatments with duplicated plates were transferred to a solid regeneration medium containing 5 mg/l of Zeatin and 150 mg/l of Timetin and allowed to incubate for 3 weeks in a light chamber at 26°C for the induction of shoots and roots from the pro-embryos of each callus. In order to obtain a concrete result, the experiment was replicated. After 3–4 weeks, the shoots and roots were regenerated. Based on the regeneration of shoots, the regeneration frequency of the callus was also measured for statistical analysis. The addition of plant hormones after autoclaving was observed to improve the induction of shoots and roots on the callus. In this case, we observed the induction of shoots and roots 10 days in advance of the expected date of shoot and root induction. In variety Navojoa, the induced juvenile leaves and shoots from the callus were shown to express GUS (Fig. 3).

**Figure 3.**
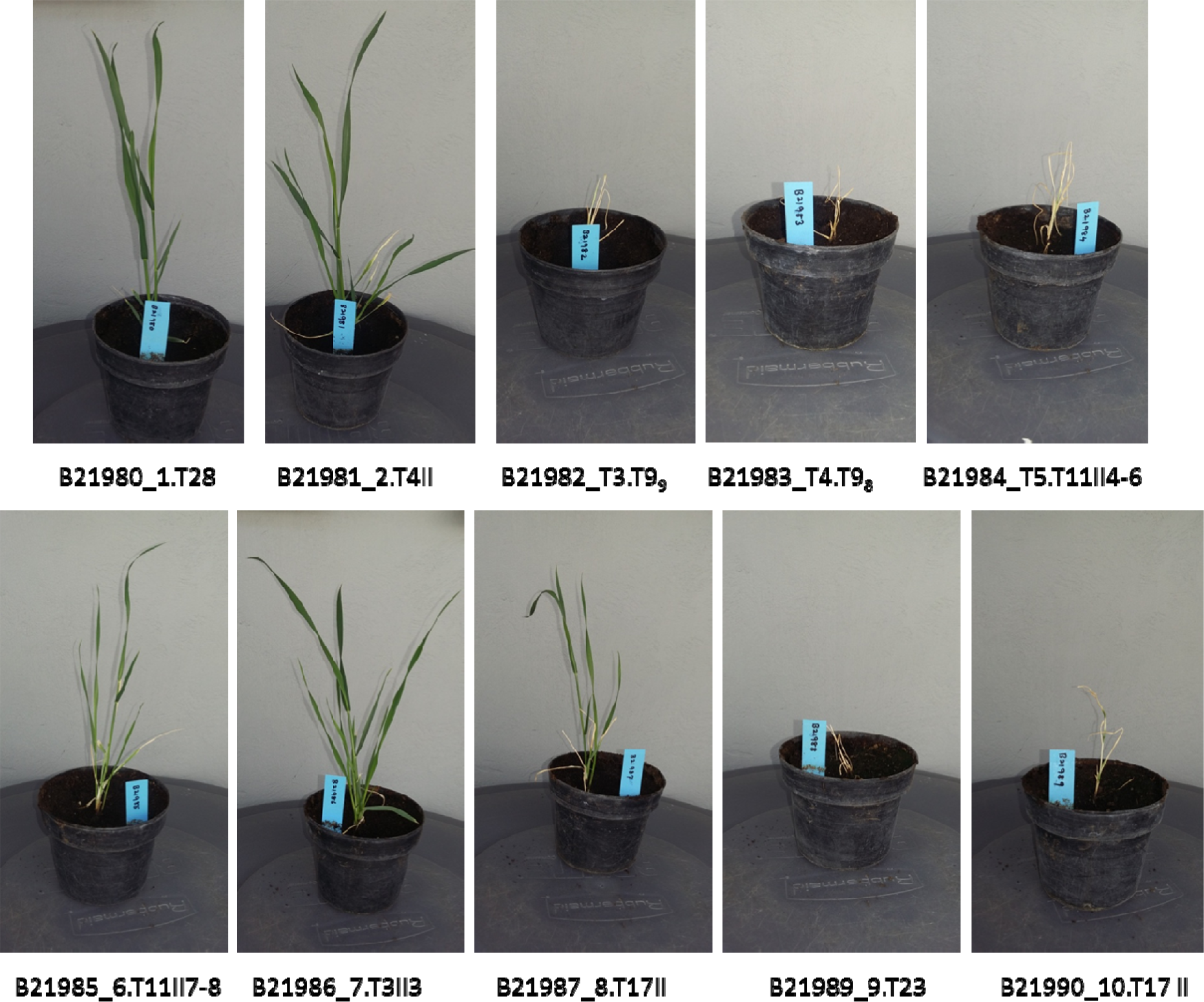
Putative transgenic Fielder genotypes subjected to hardening in a light chamber for further confirmation with PCR analysis.

### Statistical assessments for the best combinations of 2,4-D and Picloram

Different combinations of 2,4-D and Picloram resulted in different levels of phenotypic traits during the period from callus regeneration to shoot induction. However, certain treatments significantly altered the phenotypic traits.

Accordingly, the trait mean shoot frequency was higher for treatment 30 with concentrations of 48 μg/μl of 2,4-D and 64 μg of Picloram. The trait mean callus frequency for immature embryos was higher with treatments 25, 31, and 35 with concentrations of 48 μg of 2,4-D and 16 μg/μl of Picloram, 64 μg of 2,4-D and 16 μg of Picloram, and 64 μg of 2,4-D and 48 μg of Picloram, respectively. The trait mean callus size was maximum for treatment 30 with 48 μg of 2,4-D and 64 μg of Picloram, as shown in Fig. 2 and Table 1.

**Table 1.** F test statistical significance of the traits: CS: Callus size; MC: Number of regenerated Calli; NS: Number of shoots emerged from callus for the variety Fielder.

| Trait | Std. Dev | Sum Sq | Mean Sq | LSD Mean | CV | MSerror | LSD | F test | P value |
| --- | --- | --- | --- | --- | --- | --- | --- | --- | --- |
| CS | 0.82 | 33.56 | 0.96 | 6.90 | 9.32 | 0.41 | 1.31 | 0.39 | 0.00746 ** |
| MC | 2.65 | 362.4 | 10.36 | 7.22 | 26.82 | 3.75 | 3.93 | 2.80 | 0.00174 ** |
| NS | 196.37 | 196.37 | 5.61 | 3.79 | 46.62 | 3.13 | 0.85 | 1.79 | 0.0439* |

### Selection of plants against Phosphinothricin (PPT)

Shoot and root-induced calli from regeneration medium were transferred to solid 1X selection medium containing phosphinothricin (PPT) at 2.5 mg/l and 210 mg/l of timetin. These calli were allowed for the preliminary selection of transformed or transgenic plants with a marker gene called *Bar* or *pat* encoding phosphinothricin acetyl transferase, which inactivates PPT. Otherwise, in non-transgenic plants, the PPT inhibits glutamine synthetase, which leads to senescence, such that the regenerated plants would be unable to survive further because of the absence of the *Bar* gene. After phosphinothricin (PPT) selection, hardening of the plants in soil-filled pots was done timely for three weeks in a light chamber.

### Hardening of putative transgenic plants in a light chamber

The plants that showed better growth without senescence or deprivation of growth were selected from the PPT selection medium and transferred to a pot filled with fertile soil. Pot transplanted putative transgenic plants were transferred to a light chamber and allowed to grow for 2 to 3 weeks under controlled growth conditions in the light chamber (Temperature: 23.9 °C, Relative Humidity: 52%, Light illumination: 350 μMOL, CO2: 3000 ppm). During the period of light chamber-based hardening, fertilizers such as urea as a nitrogen source and phosphoric acid as a phosphate source were given at 100 mg and 10 mg quantities, respectively (Fig. 3). During the hardening process, some genotypes didn’t acclimate to grow further in the light chamber, as shown in Figure 3.

### Transferring plants from the light chamber to the green house after hardening

Pots were filled with a mixture of sand, soil, and organic matter in the ratio of 1:2:1. The green house was maintained with an experimental ambient temperature of 25.13 °C (range 23–27.8 °C) during the day and 14.29 °C (range 12–18 °C) during the night, with 8 hours of photoperiod. Similar amounts of fertilizer as described for light chamber growth were also applied. During the growth of plants, various parts were tested for GUS expression. Before attaining the reproductive stage, mature green leaves were excised and stored at -80 oC for DNA extraction. After attaining the reproductive stage, various parts of tissues were collected and used for the GUS expression assay.

### Ammonium Glufosinate or Phosphinothricin (PPT) Leaf Painting Assay

The non-transgenic genotypes showed complete necrosis of the leaves due to the absence of the *Phosphinothricin Acetyltransferase* (*Pat*)/*Bialaphos Resistance* (*Bar*) gene and the leaves being unable to neutralize the PPT, which led to complete necrosis of the leaves on the PPT-painted area. The transgenic genotypes showed a lack of necrosis or partial necrosis of the leaves due to the presence of the *Pat*/*Bar* gene, and the leaves were able to neutralize the PPT, which led to the absence of necrosis in the leaves on the painted area (Fig. 4).

**Figure 4.**
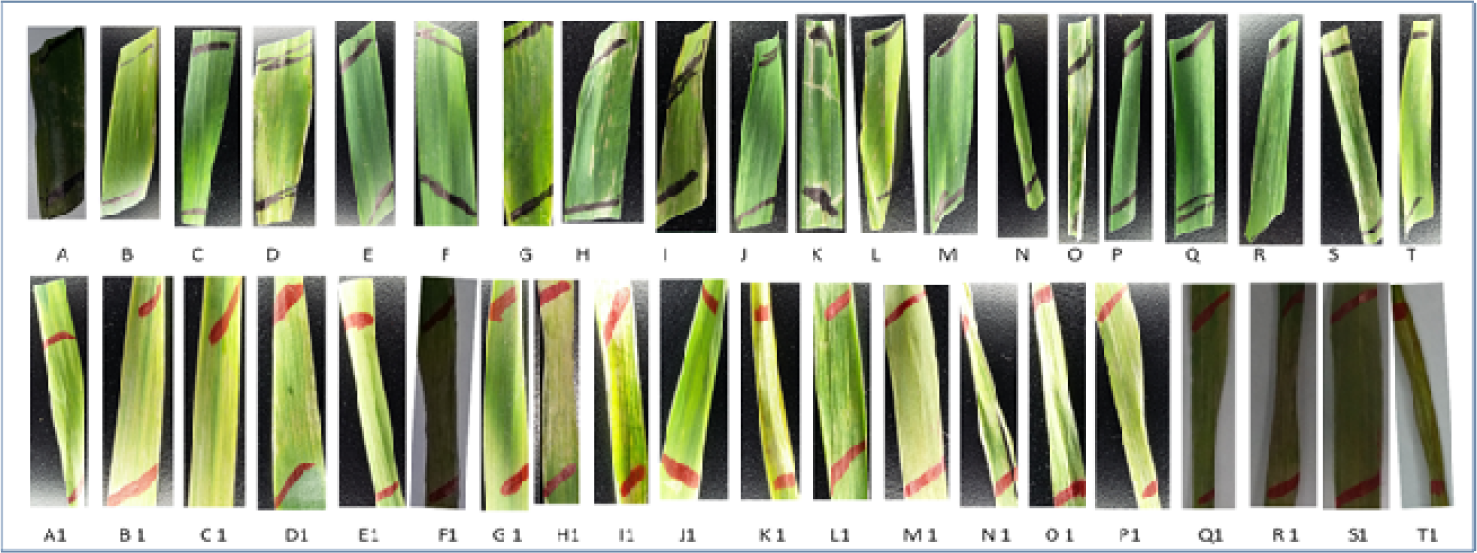
Based on the PPT leaf painting assay, putative transgenic genotypes of the wheat variety Fielder. A. 23I, B. 12II, C. 20Ib, D. 18I, E. 15IIIa, F. 25Ic, G. 15I, H. 25II, I. 12IV, J. 17IId, K. 24Ib, L. 16I, M. 25Ie, N. 14I, O. 14II, P. 15IIIc, Q. 17Ie, R. 20III, S. 20IIa. T. 14I0. Non-transgenic control genotypes of Fielder variety. A1-T1.

### *GUS* assay for various tissues from transgenic and non-transgenic genotypes

The expression of the enzyme was observed in various tissues upon the observation of blue color formation along the tissue or a specific portion of the tissue. The expression was predominated at young, developing seeds and tissues around the seed coats in various genotypes. Some of the genotypes were observed to have blue color formation in roots and anthers. The expression in leaves was much lower, and a single genotype expressed the enzyme in a very young tip of the leaf. The blue color formation was started after overnight incubation, whereas the enhanced intensity of the blue color was observed after 24 hours of incubation. The transgenic genotypes of the variety Fielder expressing the β-glucouronidase from various tissues are shown in Figure 5. Expression of β-glucouronidase was much lower in the developing leaf tissues of transgenic genotypes. The intensity of the color formation in anthers varies among genotypes, as revealed through boxes 6, 7, and 8 in Figure 5. The transgenic genotypes of the variety Fielder expressing the β-glucouronidase from various tissues are shown in Figure 5. Expression of β-glucouronidase was much lower in the leaf tissues of developing transgenic genotypes. The intensity of the color formation in anthers varies among genotypes, as revealed through boxes 6, 7, and 8, as shown in Fig. 5.

**Figure 5.**
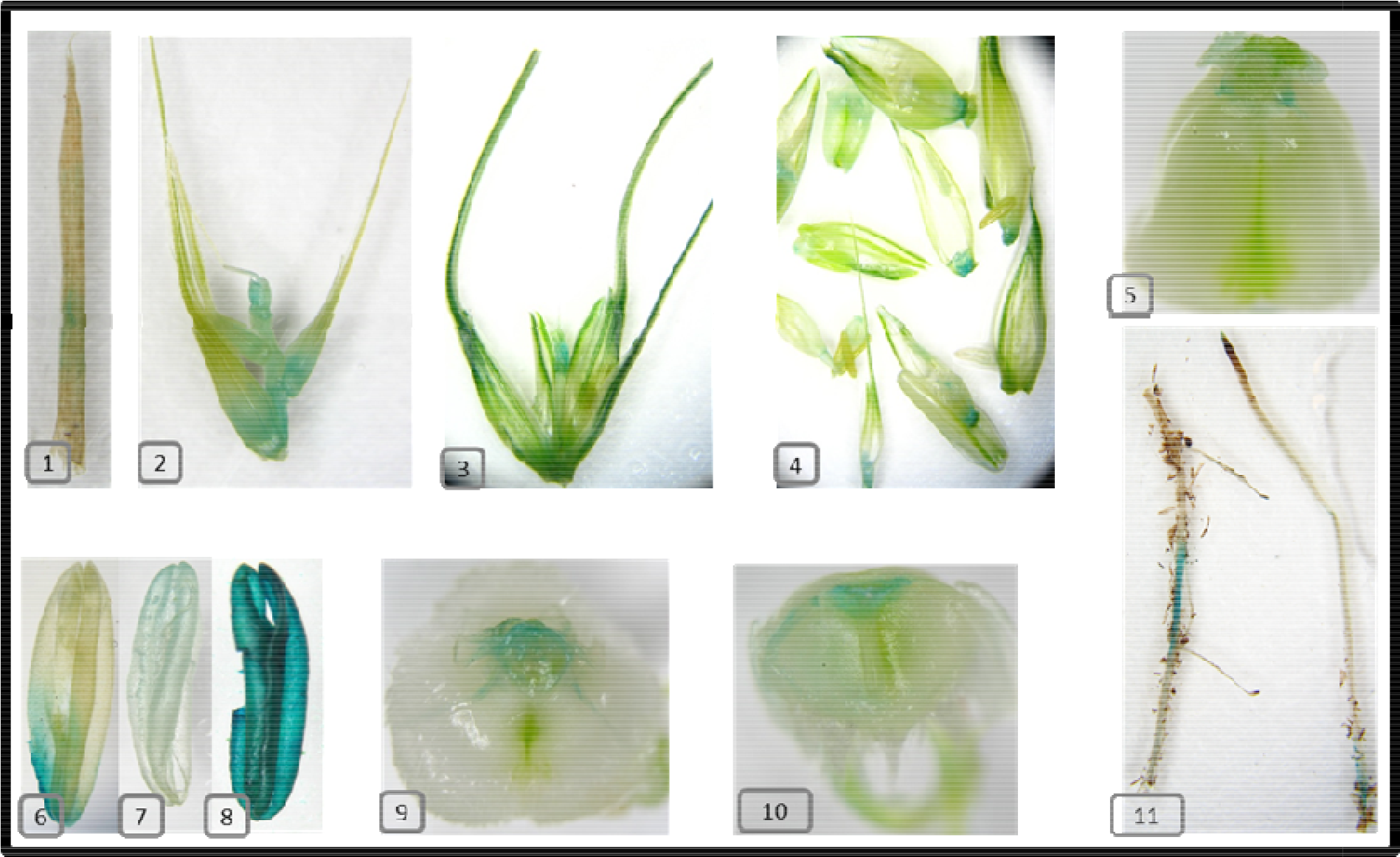
*GUS* assay from various tissues from the genotypes of the Fielder variety Samples: 1. 75I_T17, 2. 8_T17 II. 3. T4II. 4. T4II, 5. T11II4-8, 6. T28, 7. T28, 8. T23, 9. T17 II, 10. T3II, 11. T6_II_7-9.

### Further confirmation with the *Bar* gene

The same putative transgenic plants, which were used in *Bar* or *gusA* gene amplification. It is another selectable reporter gene. The expression of this gene will be detected upon using the substrate 5-bromo-4-chloro-3-indolyl glucuronide (X-Gluc). Young leaf tissues were used for its expression, and the expression of this gene was confirmed based on the observation of blue color formation along the surface of the young leaf tissue from a putative transgenic plant. The putative transgenic plants were selected based on the amplification of the *Bar* gene. The *Bar* gene is a marker gene that is used to select plants against PPT due to the expression of *Phosphinothricin* acetyltransferase, which detoxifies the PPT and enables the plant to survive. Otherwise, the plants without the *Bar* gene would not survive in PPT media. The length of the *Bar* gene inside the Ti Plasmid was ∼ 421 bp, and BAR F and BAR R primers were used to amplify the *Bar* gene from the putative transgenic plant. The amplification of the *gusA* gene is not shown here. The PCR experiment for the *Bar* gene revealed the presence of transgenic plants from randomly selected samples, which also showed better growth during PPT treatment. The result *Bar* gene fragment with the frequency of 62%, as revealed through the gel picture for the variety Baj (Fig. 6), and even up to 80% (Thiyagarajan et al., 2022).

**Figure 6.**
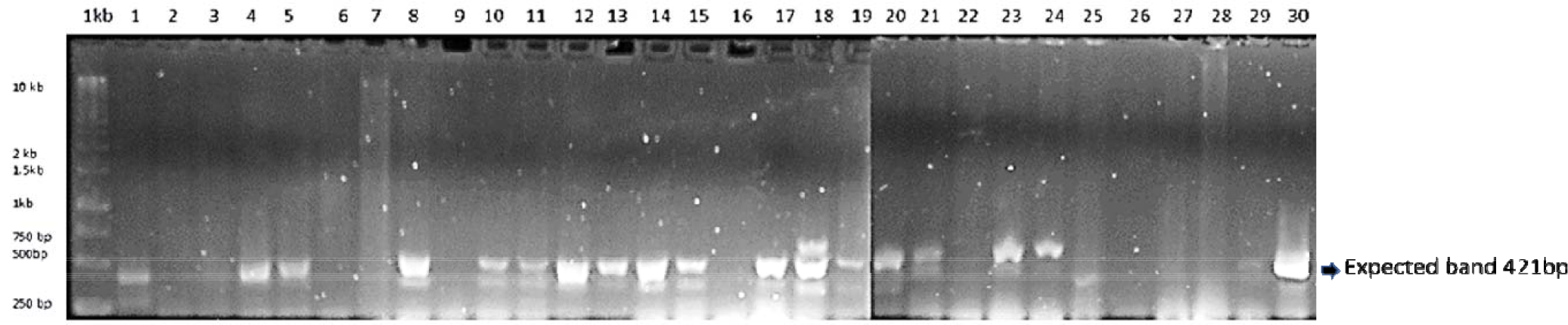
Amplification of the *bar* gene in the wheat variety Baj Genotypes: Samples: 1. 14B2, 2. 15B2, 3. 15B3, 4. 16B1, 5. 17B1, 6. 17B2, 7. 18B, 8. 19B, 9. 20B2, 10. 21B, 11. 22B1, 12. 23B1, 14. 24B, 15.24B2, 16. 25B1, 17. 25B2, 18. 29B1, 19. 29B2, 20. 30B2, 21. 30B3, 22. 31B, 23. 32B, 24. BCA1, 25. BCB2, 26. BCB1, 27. BCB3, 28. 29. 2B BCE2, 30. Plasmid Control.

## Discussion

The current study provided evidence of transgenic efficiency through *Agrobacterium*-mediated transformation in four bread wheat varieties, and extensive focus was provided for the Fielder variety with various perspectives on the transgenic analysis. The seed size is relatively smaller (5 mm) than other commercial varieties; for instance, the commercial variety of Mexico is called Navojoa, whose size was even up to 6–8 mm. Fielder seeds harvested after 14 DAF were smaller in size and thus yielded smaller embryos. Being smaller seeds, the excised embryos would be much smaller (in more circumstances, the size was <1 mM) and were completely lysed by *Agrobacterium* during the observation on callus induction medium. Therefore, embryos were collected after 14 DAF and kept in the refrigerator for 2–3 days in order to obtain 1.5 mm–2 mm size to alleviate the embryo’s lysis by *Agrobacterium* during transformation and induction.

A number of physio-chemical factors play a crucial role in the successful *Agrobacterium*-mediated transformation. Wounded tissues would naturally produce a phenolic compound called Acetosyringone to a certain extent, while the addition of 200 μM of Acetosyringone (4’-Hydroxy-3’, 5’-dimethoxyacetophenone) in both inoculation and co-cultivation medium was reportedly represented as an important signal transducer for the successful delivery of T-DNA into the host explant (Cheng et al., 1997). Likewise, another phenolic compound, so-called Pholoroglucinol (1,3,5-trihydroxybenzene), also plays a crucial role due to its synergetic effect with auxins and cytokinins (Jaime et al., 2013). For the current study, various concentrations of these substances were tested, and better efficiency of transformation and callus regeneration was obtained through the addition of 250 μM Acetosyringone and 410 μM Phloroglucinol. Besides, the effect of the surfactant Silwet-77 on increased transformation efficiency was also reported (Li et al., 2009). Thus, 0.015 μM/μl of Silwet-77 was added to the inoculation medium and shown to increase the efficiency of transformation in the inoculation medium.

Antioxidants that alleviate the effect of oxidative stresses and prevent the oxidation of phenolic compounds, including L-Cysteine and other compounds, were well characterized (Tothet al., 1994; Ziv and Halevy, 1983) during the process of *Agrobacterium*-mediated transformation. Therefore, 1 mM L-Cysteine was tested starting from 1X inoculation medium and other subsequent mediums, which were found to enhance callus regeneration and transformation efficiency upon observation of callus size and GUS expression in transformed embryos. Similarly, adsorbent PVP binds with phenolic compounds, prevents oxidation and polymerization, and further helps to prevent the activity of oxidation of phenolic compounds caused by phenolase (Sathyanarayana and Dalia B. Varghese, 2007). Accordingly, 0.5 mM PVP was included in the callus induction medium and was found to be more effective in preventing the oxidation of phenolic compounds, although the size of the callus was reduced compared to other treatments. Glycerol is a known cryoprotectant; however, it was reportedly possessed morphogenesis activity (Pilar et al., 1998), and exogenous application has a potential effect on plant growth (Eastmond, 2004). Thus, 1% glycerol treatment during the transformation of *Agrobacterium* in 1X liquid inoculation medium enhanced the transformation efficiency based on the observation of a higher number of blue spots on embryos from the X-Gluc buffer treatment.

Co-cultivation media was supplemented with 20 mg/L of 2,4-D and 20 mg/L of Picloram and reports suggested that adding these kinds of hormones can suppress the precocious germination of immature embryos (Jones et al., 2005) during the early phase after the transformation and co-cultivation, while some reports suggested that the removal of the root axis would prevent zygotic germination (Weir et al., 2001). External hormonal supplements are crucial for plants that would naturally produce limited auxins (Mendoza et al., 2002). In tissue culture, the supplementation of 2,4-D and other hormones would cause callus formation and regeneration and, in some cases, even somoclonal variation, so it is mandatory to optimize the level of necessity during callus regeneration through *Agrobacterium*-mediated transformation. Therefore, for the current study, 36 different combinations of 2,4-D and Picloram were tested for callus regeneration efficiency in callus induction medium. Some of the combinations had a significant effect on the regeneration of calluses based on the observation of callus frequency, callus size, and shoot and root regeneration frequencies. The mean callus size was minimum with a range of 5.875 mm for the combination of 16 μg of 2,4-D and 16 μg of Picloram (treatment 1) and 16 μg of 2,4-D and 16μg of Picloram (treatment 36), while maximum with a range of 7.175mm for treatment 30 with the combination of 48 μg of 2,4-D and 64 μg of Picloram. After that, the callus regeneration percentage was lesser (37.5%) for treatment 20 with the combination of 40μg of 2,4-D and 24 μg of Picloram, while it achieved a higher number (97.5%) for treatments 25, 31, and 35 with the combination of 48 μg of 2,4-D and 16 μg of Picloram, 64 μg of 2,4-D and 16 μg of Picloram and 64 μg of 2,4-D and 48 μg of Picloram, respectively. Likewise, the mean shoot regeneration frequency was 10% for treatment 20, which showed less with 40 μg of 2,4-D and 24 μg of Picloram, while treatment 30 attained maximum (57.5%) with a combination of 48 μg of 2,4-D and 64 μg of Picloram.

From the regeneration media, individual shootlets were transferred to Phosphinothricin (PPT) media and allowed to grow for three weeks to obtain PPT-resistant plants. The plants, which showed better growth and greenish color, were presumed to be putatively transgenic and were transferred to a light chamber and subsequently to a green house. Even after optimization, *bar* gene primers amplified gene-specific fragments and other non-specific fragments, while *gusA* gene primers selectively amplified GUS-specific fragments for confirmation of the transgenic nature of the genotypes. It was reported that the incomplete integration of T-DNA would bear either the *bar* or the *gusA* gene (Wu et al., 2006). Likewise, the expression of the *gusA* gene is restricted to certain genotypes, and in others, suppression of GUS expression occurs due to gene silencing, incomplete integration of T-DNA, or rearrangements. Transgenic confirmation with *bar* gene amplification in Baj and Kachu varieties and the segregation pattern of transgenic progenies (seed-based analysis) from the variety fielder were reported in our recently published preprint (Thiyagarajan et al., 2022). After *gusA*-PCR confirmation, various parts of the plants were subjected to GUS expression analysis and showed the formation of a blue color around the tissues due to the expression of the *gusA* gene, and some of the *gusA-*PCR plants have not shown GUS expression, which is presumed due to the absence or an inactive form of the *gusA* gene.

## Conclusion

The current study revealed that moderate to higher efficiency of transformation in wheat was achieved with appropriate modifications and the inclusion of specific compounds at appropriate stages in growth media. Secondly, apart from the nutritional factors, the physical factors were also playing a vital role, which induced stress on the *Agrobacterium* to enhance the transformation efficiently. The current study was focused on optimizing both of these major factors. The results obtained from this study through appropriate modifications of existing protocols can be effectively used for *Agrobacterium*-mediated transformation, even though the transformation efficiency may vary from variety to variety according to the results obtained from this study in accordance with previous reports based on the genetic structure, expression pattern, and coordination of genes of a specific genotype or variety. This protocol optimization article aims to improve transgenic efficiency across selected different varieties; hence, each variety or accession may require an appropriate modification of a protocol based on its physiological features, genetic structure, and epigenetic mechanism.

## Authors Contribution

Wrote the manuscript: KT; Wheat tissue/embryo culture, *Agrobacterium* transformation: LN, KT, MP; Statistical analysis, transgenic confirmation through biochemical and molecular biological aspects: KT; Suggestions for improving manuscript; GV, PV.

## Acknowledgement

Authors duly acknowledge Dr. Kanwarpal Singh Dhugga (Head and Principal Scientist, Biotechnology, CIMMYT) for his valuable guidance to carry out this research study. Authors also grateful to Dr. Kevin Pixley (Deputy Director General for Research (Breeding and Genetics), a.i., and Director of the Genetic Resources Program, CIMMYT) and Ms. Rodelita Panergalin (Program Manager: Genetics Resources Program, CIMMYT) for their encouragement and supports.

## References

1. Jaime A. Teixeira da Silva, Judit Dobránszki, Silvia Ross. In Vitro Cell.Dev.Biol.-Plant (2013) 49: 1.

2. Cheng M, Fry JE, Pang SZ, Zhou HP, Hironaka CM, Duncan DR, Conner TW, Wan YC. Genetic transformation of wheat mediated by Agrobacterium tumefaciens. Plant Physiology. 1997;115:971–980.

3. FAO Cereal Supply and Demand Brief, 2023 (https://www.fao.org/worldfoodsituation/csdb/en/)

4. Jian-Feng Li, Eunsook Park, Albrecht G von Arnim and Andreas Nebenführ,The FAST technique: a simplified Agrobacterium-based transformation method for transient gene expression analysis in seedlings of Arabidopsis and other plant species. Plant Methods 2009 5:6.

5. Toth K., Haapala T., Hohtola A. Alleviation of browning in oak explants by chemical pretreatments. Biol. Plant. 1994;36:511–517.

6. Ziv, M.; Halevy, A. H. Control of oxidative browning and in vitro propagation of Strelitziareginae. HortScience 18(4):434–436;1983.

7. Eastmond PJ (2004) Glycerol-insensitive Arabidopsis mutants: gli1 seedlings lack glycerol kinase, accumulate glycerol and are more resistant to abiotic stress. Plant J 37: 617–625.

8. García Jiménez, Pilar; Rodrigo, Marta; Robaina, Rafael. Influence of plant growth regulators, polyamines and glycerol interaction on growth and morphogenesis of carposporelings of Grateloupia cultured in vitro. Journal of Applied Phycology, 1998, vol. 10, p. 95–100.

9. Plant Tissue Culture: Practices and New Experimental Protocols, B. N. Sathyanarayana and Dalia B. Varghese, I.K. International Publishing House Pvt. limited, 2007, ISBN 81-89866, 11–7.

10. Jones HD, Doherty A, Wu H. Review of methodologies and a protocol for the Agrobacterium-mediated transformation of wheat. Plant Methods. 2005;1:5.

11. Weir B, Gu X, Wang MB, Upadhyaya N, Elliott AR, Brettell RIS.Agrobacteriumtumefaciensmediated transformation of wheat using suspension cells as a model system and green fluorescent protein as a visual marker. Aust J Plant Physiol. 2001;28:807–818.

12. Mendoza MG, Kaeppler HF. Auxin and sugar effects on callus induction and plant regeneration frequencies from mature embryo of wheat (Triticum aestivum L.) In Vitro Cell Dev Biol. 2002;38:39–45.

13. Wu H, Sparks CA, Jones HD. Characterisation of T-DNA loci and vector backbone sequences in transgenic wheat produced by Agrobacterium-mediated transformation. Molecular Breeding.2006;18:195–208.

14. Lazo GR, Stein PA, Ludwig RA. A DNA transformation-competent Arabidopsis genomic library in Agrobacterium. Bio-Technology 1991;9:963–967.

15. Luo J, Li S, Xu J, Yan L, Ma Y, Xia L. Pyramiding favorable alleles in an elite wheat variety in one generation by CRISPR-Cas9-mediated multiplex gene editing. Mol Plant. 2021 Jun 7;14(6):847–850.

16. Hellens RP, Edwards EA, Leyland NR, Bean S, Mullineaux PM. pGreen: a versatile and flexible binary Ti vector for Agrobacterium-mediated plant transformation. Plant Molecular Biology 2000;42:819–832.

17. Bourdon V, Harvey A, Lonsdale DM. Introns and their positions affect the translational activity of mRNA in plant cells. EMBO Reports 2001;2:394–398.

18. Ke XY, McCormac AC, Harvey A, Lonsdale D, Chen DF, Elliott MC. Manipulation of discriminatory T-DNA delivery by Agrobacterium into cells of immature embryos of barley and wheat. Euphytica 2002;126:333–343.

19. Christensen AH, Quail PH. Ubiquitin promoter-based vectors for high-level expression of selectable and/or screenable marker genes in monocotyledonous plants. Transgenic Research 1996;5:213–218.

20. R Core Team. 2014, R: A language and environment for statistical computing. R Foundation for Statistical Computing, Vienna, Austria. URL http://www.R-project.org/.

21. Ming Cheng, Joyce E. Fry, Shengzhi Pang, Huaping Zhou, Catherine M. Hironaka, David R. Duncan, Timothy W. Conner, and Yuechun Wan, Genetic transformation of wheat mediated by Agrobacterium tumefaciens. Plant Physiol., 1997, 115, 971–980.

22. Ozawa K, A high-efficiency Agrobacterium-mediated transformation system of rice (Oryza sativa L.), Methods Mol Biol. 2012; 847:51–7.

23. Ishida Y 1, Hiei Y, Komari T, Agrobacterium-mediated transformation of maize. Nat Protoc. 2007;2(7):1614–21.

24. Y. He, H. D. Jones, S. Chen, X. M. Chen, D. W. Wang, K. X. Li, D. S. Wang and L. Q. Xia, Agrobacterium-mediated transformation of durum wheat (Triticum turgidum L. var. durum cv Stewart) with improved efficiency, J. Exp. Bot. (2010) 61 (6): 1567–1581.

25. Songstad DD, Somers DA, Griesbach RJ (1995) Advances in alternative DNA delivery techniques. Plant Cell Tiss Org Cult, 40: 1–15.

26. Ishida Y, Tsunashima M, Hiei Y, Komari T, Wheat (Triticum aestivum L.) transformation using immature embryos, Methods Mol Biol. 2015;1223:189–98.

27. Terese Richardson, Jenny Thistleton, T. J. Higgins, Crispin Howitt, Michael Ayliffe, Efficient Agrobacterium transformation of elite wheat germplasm without selection, Plant Cell Tiss Organ Cult (2014) 119: 647.

28. Bartlett JG, Alves SC, Smedley M, Snape JW, Harwood W. High-throughput Agrobacterium-mediated barley transformation. Plant Methods 2008; 22:1–12.

29. Jones, H.D. (2005). Wheat transformation: current technology and applications to grain development and composition. Journal of Cereal Science, 41, 137–147.

30. Amoah, B.K.; Wu, H.; Sparks, C. & Jones, H.D. (2001). Factors influencing Agrobacterium mediated transient expression of uidA in wheat inflorescence tissue. Journal of Experimental Botany, 52, 1135–1142.

31. Ke, X.Y, McCormac, A.C, Harvey, A, Lonsdale, D, Chen, D.F, Elliott, M.C. (2002). Manipulation of discriminatory T-DNA delivery by Agrobacterium into cells of immature embryos of barley and wheat. Euphytica, 126, 333–343.

32. Wu, H, Doherty, A. & Jones, H.D. (2008). Efficient and rapid Agrobacterium-mediated genetic transformation of durum wheat (Triticum turgidum L. Var. Durum) using additional virulence genes. Transgenic research, 17, 425–436.

33. He, Y, Jones, H.D.; Chen, S, Chen, X.M, Wang, D.W, Li, K.X, Wang, D.S, Xia, L.Q. (2010). Agrobacterium-mediated transformation of durum wheat (Triticum turgidum L. Var. Durum Cv Stewart) with improved efficiency. Journal of Experimental Botany, 61, 1567–1581.

34. Wu H, Sparks CA, Amoah B, Jones HD. Factors influencing successful Agrobacterium-mediated genetic transformation of wheat. Plant Cell Reports. 2003;21:659–668.

35. Stachel S. E., Messens E., van Montagu M., Zambryski P. Nature. 1985;318:624–629.

36. Mónica Blanco, Roberto Valverde 2, Luis Gómez, Optimization of genetic transformation with Agrobacterium rhizogenes, Agronomía Costarricense 27(1): 19–28. 2003.

37. Wernicke W, Milkovitz L. Effect of auxin on mitotic cell cycle in cultured leaf segments at different stages of development in wheat. Plant Physiology. 1987;69:16–22.

38. Möller B, Weijers D. Auxin control of embryo patterning. Cold Spring Harb Perspect Biol. 2009;1.

39. Thiyagarajan K, Noguera LM, Pacheco M, Govindan V, Vikram P, Agrobacterium mediated transformation and deciphering SNPs in TaLr67 gene homeologs for gene editing in wheat, bioRxiv 2022.03.23.485492.

40. Tilman D, Balzer C, Hill J, Befort BL (2011) Global food demand and the sustainable intensification of agriculture. Proc Natl Acad Sci USA 108, 20260–20264.

